# Targeted engineering of the phase separating PARCL protein

**DOI:** 10.1101/2024.06.21.600009

**Authors:** Ruth Veevers, Steffen Ostendorp, Anna Ostendorp, Julia Kehr, Richard J. Morris

## Abstract

PARCL is a plant-specific RNA-binding protein (RBP) that exhibits chaperone activity, is abundant in the phloem, intrinsically disordered, and contains a prion-like domain (PLD). PARCL proteins have been observed to form large biomolecular condensates *in vivo* and *in vitro*. Biomolecular condensates are membraneless compartments, wherein biomolecules become partitioned from their surrounding liquid environment into liquid droplets with their own composition, dynamics, and function. Which molecular properties drive phase separation is of great interest for targeted engineering efforts. Here, we present results on residue interactions derived from simulations of PARCL using course-grained molecular dynamics with the HPS-Urry model. We adjust the parameters of the simulations to allow the inclusion of folded eYFP tags, since fluorescent tags are often used in phase separation experiments for visualising droplets, yet have not been included in simulations to date. While still simulating phase separation, these trajectories suggest minor changes to droplet and network structure when proteins contain eYFP. By analysing the residues of the PARCL molecules that come within contact distance in the simulations, we identify which individual residues drive phase separation. To experimentally validate these findings, we introduced mutations of the most contacted residues and could indeed confirm that these mutations prevent the formation of condensate droplets. To investigate the RNA-binding of PARCL, we added microRNA to the simulation and find a short region of PARCL consistently making contact with the miRNA, which is also in agreement with predictions and experiments. We discuss the implications of our findings in terms of model-guided engineering of biomolecular condensates.

## Introduction

Biomolecular condensates are membraneless compartments, such as stress granules and nucleoli, which can perform key biological functions (Hirose et al, 2023; Protter and Parker, 2016; Toretsky and Wright, 2014). These condensates are often formed by the process of liquid-liquid phase separation (LLPS), wherein biomolecules become partitioned from their surrounding liquid environment into liquid droplets with their own composition and dynamics (Brangwynne et al, 2009; Hyman et al, 2014). From a thermodynamic viewpoint, we would expect condensate formation to be sensitive to macroscopic factors, such as temperature, pressure, and concentration, and these dependencies have been experimentally demonstrated, leading to the determination of phase diagrams for different specific proteins (Hyman et al, 2014, Li et al, 2021, Milkovic and Mittag, 2020). From a molecular viewpoint, the balance between intra- and intermolecular forces and indeed relationships between molecular composition and phase separation have been discovered (Saar et al, 2021; Ruff et al, 2022). For instance, intrinsic disorder and the presence of a prion-like domain (PLD) are common to phase separating proteins (Martin and Holehouse, 2020; Wang et al, 2018). Amino acid properties, such as hydrodynamic size, propensity for beta-turns, and sequence hydrophobicity, also influence condensate formation and the amino acid distribution differs significantly between disordered proteins that are prone to phase separation and those that are not (Ibrahim et al, 2022). Multivalent interactions drive condensate formation, and the types and patterning of proteins’ amino acids can suggest which interactions are possible for a protein. The contributions of electrostatic interactions are dictated by charged residues and how they are distributed within the protein sequence (Das and Pappu, 2013). Aromatic residues are a strong determinant of phase separation behaviour (Martin et al, 2020) due to their contribution to 𝜋-𝜋 and cation-𝜋 interactions (Vernon et al, 2018).

Key insights into phase separation have been gained from computational studies, with approaches ranging from 1D sequence analyses (‘low resolution’), such as the method of residue-counting proposed by Wang et al (2018), to spatio-temporal analyses, such as molecular dynamics (MD) simulations that calculate the trajectories of individual atoms (‘high resolution’) (Zheng et al, 2020). Whilst recent advances in molecular dynamics are rapidly expanding their range of applicability (Antolinez et al, 2024, Phillips et al, 2020), studying large systems over long timescales using all-atom MD simulations can quickly become computationally prohibitive. To address this, a common approximation is to build coarse-grained models where groups of atoms are represented by a single particle or “bead” (Martin et al, 2020; Ruff, et al, 2015; Marrink et al, 2007; Monticelli et al, 2008; Tsanai et al, 2021; Joseph et al, 2021). The bead and interaction parameters are determined from experimental data and by the outcomes of smaller all-atom simulations. Dignon et al (2018) proposed two coarse-grained, one-bead-per-residue models in a slab simulation framework: the KH model is based on the Miyazawa-Jernigen potential for inter-residue attractions; the HPS model is based on a hydrophobicity scale (Kapcha and Rossky, 2014). This level of approximation allows for residue-specific effects to be taken into account. Schuster et al (2020) used coarse-grained MD to simulate the disordered domain of the LAF-1 protein, finding that a short, conserved, hydrophobic region was driving phase separation. To assess the relative effects of electrostatic, hydrophobic, 𝜋-𝜋 and cation-𝜋 interactions, Das et al (2020) produced simulations using the models proposed in Dignon et al (2018), both as published and tailored to include cation-𝜋 interactions. They simulated the IDR of the Ddx4 protein and three variants with less propensity to phase separate, identifying areas of agreement and disagreement between simulations produced by the HPS model and phase separation behaviour ascertained experimentally. Regy et al (2021) improved the Dignon et al (2018) HPS model’s agreement with experimental observations by replacing the Kapcha-Rossky hydrophobicity scale with that of Urry et al (1992).

Based on advances towards developing a molecular grammar that explains phase separation, Kilgore and Young (2022) anticipated a complete grammar that would clarify the relationships between amino acid sequence features and phase separation propensity. Molecules could thus be designed that could be specifically recruited into a condensate, or that could modulate the phase behaviour or material properties in a system. Many studies have attempted to find mutations that would change an existing protein’s LLPS propensities (Bracha et al, 2019, Ji et al, 2022). These mutations were proposed using various methods: predictive screening of exhaustive random mutations (Bolognesi et al, 2019), patient-derived mutations (Patel et al, 2015), or *in silico* homology modelling (Du et al, 2019). Coarse-grained MD simulations have been used to find short regions to delete based on contact probability (Schuster et al, 2020), to guide the progress of genetic algorithms (Lichtinger et al, 2021), and to form the basis of shorter or smaller all-atom simulations (Zheng et al, 2020).

In this work, we use the coarse-grained HPS-Urry model (Regy et al, 2021) to simulate the phase separating PARCL protein. Higher resolution simulations have been used to make confident assessments of the specific interactions driving LLPS (Zheng et al, 2020, Schuster et al, 2020). However, this is a more computationally expensive process, whereas the speed and ease of the coarse-grained model offers the possibility of high-throughput data generation that may provide sufficient insights to guide targeted engineering. The target protein, PARCL, is an RNA-binding protein (RBP) recently characterised (Ostendorp et al, 2022) as a phloem-abundant, intrinsically disordered protein that exhibits chaperone activity and contains a PLD. PARCL proteins form large condensates observed *in vivo* and *in vitro*, highlighted with the addition of using fluorescent eYFP domains. Phase separation simulations have not previously included fluorescent tags which are often used in phase separation experiments for visualising droplets, especially *in vivo*. The presence and choice of fluorescent tags can affect LLPS (Uebel and Phillips, 2019; Alberti et al, 2018). Therefore, we adjusted the parameters of the simulations to allow the inclusion of folded eYFP tags, which enabled us to identify the effects of eYFP on PARCL’s phase separation behaviour. The inclusion of eYFP still led to phase separation in the simulation, however, their trajectories suggest changes to droplet and network structure may occur when proteins contain eYFP. We score individual residues using a radius-based contact distance estimation to predict which interactions may drive phase separation, and validate these findings experimentally by mutation of the most highly-scoring residues. We demonstrate that inducing the mutations predicted by the simulations can indeed prevent the formation of condensate droplets *in vitr*o and in plant leaves *in vivo*. Adding microRNA to the system allows us to recapitulate PARCL’s experimentally determined RNA-binding behaviour both in the binding location and in the observed effect on condensates. We therefore demonstrate that residue-level approximations are sufficient to guide targeted engineering that influences phase separation and RNA binding of PARCL and, potentially, other proteins.

## Methods

### Framework

Fig 1 shows the steps we took to produce and analyse the simulated MD trajectories of multiple interacting units of the same protein, starting from a given protein sequence. We further explain these steps in the following sections.

**Figure 1.**
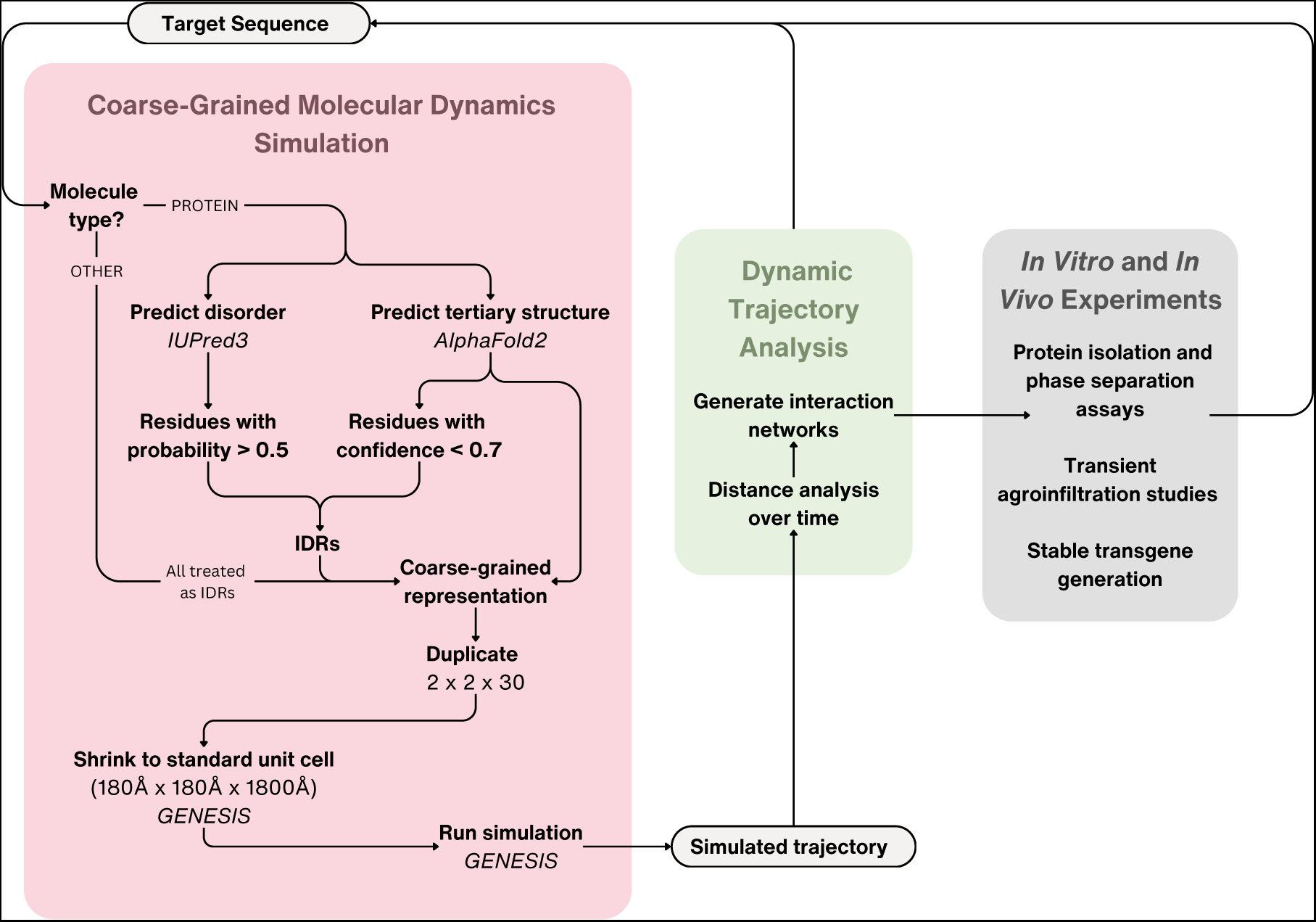
Representation of the framework used to generate molecular dynamics simulations from a sequence, analyse the resulting trajectories, and propose mutations to disrupt phase separation.

The initial simulation set-up differs based on the type of the target sequence. For proteins, 3D coordinates for the tertiary structure are predicted using AlphaFold2 (Jumper et al, 2021) and regions of disorder are predicted with IUPred3 (Erdős, Pajkos, and Dosztányi 2021). The IUPred3 output and the AlphaFold2 confidence scores are used to establish which parts of the structure should be treated as folded and which parts should be treated as IDRs. From the predicted structure and predicted list of IDRs, a coarse-grained representation of the protein is generated using a GENESIS tool. Other molecule types are all treated as fully intrinsically disordered, and CG representations are generated directly from the sequence as a line of beads which is relaxed with a short period of simulation. A further GENESIS tool is used to duplicate the molecules into a grid, and two GENESIS simulations are run. Each establish a “unit cell” surrounding the grid of molecules and imposing periodic boundary conditions such that if a bead leaves the cell boundaries, it re-enters with the same trajectory from the opposite boundary. This simulates a system of infinite size containing infinite repetitions of the contents of the central unit cell. In the first simulation the bounds of this unit cell are placed around the grid of molecules and then gradually shrunk until the unit cell reaches a uniform target size. The second, longer simulation takes the output of the shrinking simulation and runs for 100 nanoseconds to capture phase behaviour. The dynamic coordinates at intervals along the simulated trajectory are then analysed to find distances between pairs of atoms, taking note of pairs of residues that come within the sum of their radii. These are used to identify attraction hubs to mutate in further simulations and in *in vitro* experiments.

### Preprocessing and set-up of initial structures

As proteins can have both structured and disordered regions, we use two external tools to derive structural information. IUPred3 (Erdős, Pajkos, and Dosztányi 2021) is strictly a predictor of disorder, which we run in its short disorder mode with medium smoothing to obtain a list of regions within the sequence that are predicted to be disordered.AlphaFold2 (Jumper et al, 2021) gives a prediction of the structure of ordered regions and assigns a confidence score to its prediction for each residue. Residues assigned low confidence by AlphaFold2 are added to the regions of predicted disorder for two reasons: it has been shown to be correlated with disorder (Tunyasuvunakool et al, 2021; Binder et al, 2022; Aderinwale et al, 2022) and it prevents over-reliance on structures that are likely to be incorrect.

### Building and running CG MD simulations

The following steps are carried out within GENESIS (J. Jung et al, 2015). We use a coarse-grained slab simulation for phase separation as described in Regy et al (2021), which builds on Dignon et al (2018) and Schuster et al (2020).

We create a coarse-grained representation of the predicted structure which uses one “bead” particle to represent each residue, centred on its C_alpha_ atom. Input files created alongside this representation specify parameters such as native contacts and angles. To these files, we add markers denoting the locations of predicted IDRs.

We duplicate this representation of the molecule, creating identical copies along the 𝑥, 𝑦, 𝑧 axes to create a 2x2x30 grid of molecules. The dimensions of this system will vary, and so to compare behaviour at similar concentrations we perform a shrinking step to unify the final volume. This step is a short simulation using the periodic boundary condition in which we reduce the unit cell dimensions to 180 Å by 180 Å by 1800 Å. This results in a protein concentration of 3.4 mM which is much higher than the 10 µM used *in vitro*, so we do not add a crowding agent.

For the second simulation, we use the periodic boundary condition with a fixed unit cell of 180 Å by 180 Å by 1800 Å and an implicit solvent with an ionic strength of 0.15 M. The volume and the temperature, 300 K, are held constant throughout the simulation. As in Regy et al (2021), the hydrophobicity scale parameters for amino acids are taken from Urry et al (1992), and the parameters for nucleic acids are taken from Regy et al (2020). We run each simulation for 10,000,000 steps, with each step corresponding to 0.01 picoseconds. The coordinates of all beads in the system are written out to a coordinate file every 10,000 steps, which results in a 1,000-frame trajectory.

Inclusion of eYFP tagging requires some adjustment of parameters. The folded eYFP domain is a similar size to the disordered proteins on which it is attached, so there are large rigid sections up to around 80 Å which obstruct intermolecular attractions that would induce phase separation. We extend the cut-off distances used by the CG model to 60 Å, the maximum for a unit cell of our standard size and raise the model’s force constant “epsilon” parameter that weights hydrophobicity interactions from the default 0.2 to 0.25. This allows these longer-range interactions to be taken into account. We used the original parameter set and the adapted eYFP parameter set to simulate the same system in which wild-type PARCL and free eYFP proteins were both present to confirm that PARCL’s overall LLPS behaviour was not changed.

### Dynamic analysis of LLPS simulations

We simulate at a coarse, residue-level resolution so it is not possible to assess exact interatomic interactions. Instead, we focus on residues that pass one another within a close enough distance to potentially make contact. We consider two residues to be in potential contact if the distance between them is less than the sum of their radii calculated from estimated residue volumes from Zamyatnin (1972).

Trajectory images are rendered in VMD (Humphrey, Dalke, and Schulten, 1996) using Tachyon ray-tracing.

### *In vitro* phase separation assays

EYFP-tagged and untagged PARCL proteins were produced and purified as described previously (Ostendorp et al. 2022) and set to 200 µM as stock concentration. To induce protein condensation, purified proteins were diluted to a final concentration of 10 µM with 1x condensation buffer (50 mM Tris-HCl pH 7.5, 150 mM NaCl). Subsequently, 10% (w/v) PEG3350 was added as a crowding agent and incubated for 2 mins prior to the measurements. 5 µl were transferred onto a glass slide (Labsolute, Th. Geyer GmbH, Germany) and carefully covered with a siliconised circular cover slide (22 mm diameter, Jena Bioscience, Germany). Condensation formation was observed by fluorescence microscopy using a BX-810 fluorescence microscope (Keyence, Germany) using a 100x oil immersion objective equipped with an eYFP filter system (Chroma, USA). All microscopic analyses were performed at an exposure time of 17 ms as a single z-plane without any black balance.

### Sampling eYFP accessibility in PARCL condensates

To test if eYFP is solvent-exposed and thus accessible within PARCL condensates, studies with an anti-YFP-nanobody (Addgene plasmid #61838) (Hartano et al. 2015, PubMed 25964651) were carried out. 15 µl glutathione high capacity magnetic agarose beads slurry (Millipore, Germany) were transferred to 1.5 ml reaction tubes and washed 3 times with 200 µl condensation buffer. Beads were coupled by incubating with 100 µM anti-GFP-nanobodies in condensation buffer for 30 minutes at RT with shaking at 150 rpm. Free nanobodies were removed by three wash steps with 200 µl condensation buffer. Then 20 µM eYFP or wildtype eYFP-AtPARCL in condensation buffer, which was supplemented with 10 % PEG3350 and incubated for 20 minutes, was directly transferred to the beads. After 30 minutes of incubation, samples were transferred onto a microscope glass slide, covered with a cover slip and observed by fluorescence microscopy using the Keyence BZ-X810 fluorescence microscope equipped with an eYFP filter.

### *In vivo* condensation studies

Agrobacteria (LBA4404) were cultivated on solid LB medium (supplemented with 100 µg/ml Kanamycin) at 28 °C overnight. Bacterial colonies were resuspended in infiltration medium (1/4 MS, 100 µM acetosyringone, 1 % sucrose, 0,005% Silwet 77 pH 6.0) and diluted to an OD600 of 0.5. These suspensions were infiltrated into 6 week old tobacco plant leaves. After cultivation of the plants over night at 25 °C in dark, plants were cultivated at 25 °C under low light conditions for further two to three days prior to sampling. Infiltrated areas were cut out, mounted onto a glass slide and covered with a glass cover slip. Cells were microscopically analysed using a Keyence BZ-X810 equipped with an eYFP filter.

## Results

### Simulation of eYFP-PARCL proteins displays phase separation behaviour

Most *in silico* simulations omit well-folded domains like fluorescence tags that are, however, often added in experiments to visualise LLPS. This might limit the predictive power of those simulations for *in vitro* and *in vivo* observations. To predict which residues in PARCL might be responsible for condensate formation, we first sought to reproduce the experimentally observed phase separation *in silico*. The PARCL proteins used for confocal microscopy in Ostendorp et al (2022) were tagged with eYFP to make the resulting droplets visible. We therefore asked whether eYFP-PARCL would undergo phase separation *in silico*. We used a coarse-grained molecular dynamics approach (see Methods) to simulate how eYFP-PARCL proteins move and interact over time (trajectories). The eYFP domain is structurally ordered unlike the mostly disordered PARCL protein. This introduces complications for MD in that different force-field parameters are required to maintain the folded protein structure of eYFP during the simulations. We used AlphaFold2 to predict the eYFP-PARCL 3D structure, which produced both an estimated set of coordinates and a confidence score for each residue. Parts with a confidence score below 0.7, or a disorder prediction above 0.5 using IUPred3, were considered unfolded and treated with the intrinsically disordered hydrophobicity force-field for phase separation. The high confidence parts (folded) were handled with a standard force-field (see Methods). Starting from a uniform distribution, we investigated whether phase separation occurs. We simulated the trajectories of 120 eYFP-PARCL proteins across 100 nanoseconds and stored their positions at 1,000 regularly-spaced time intervals for analysis and visualisation.

Fig 2A shows the final set of coordinates, or frame, where the proteins have formed droplets. The formation of eYFP-PARCL clusters as a consequence of their intermolecular interactions is consistent with phase separation. The proteins form droplets, but interestingly the eYFP domains remain at the interface rather than fully mixing with the other molecules. Given the complexity of setting up different forcefields for different parts of the protein, we considered that this mode of droplet formation was likely the result of the effects of the chosen model and forcefields, and thus an artefact. To test this idea, we used anti-GFP nanobody-coupled magnetic agarose beads to test the accessibility of the eYFP in the condensed states of eYFP-PARCL. We reasoned that nanobodies would not bind unless the eYFP was exposed. Surprisingly, these experiments revealed binding of the nanobodies to the condensates (Fig 2B), suggesting that eYFP is indeed accessible and demonstrating consistency with the simulation results. In contrast to free eYFP that showed an even distribution when bound on magnetic beads, eYFP-PARCL clearly formed liquid-like clusters on the beads appearing as bright patches.

**Figure 2.**
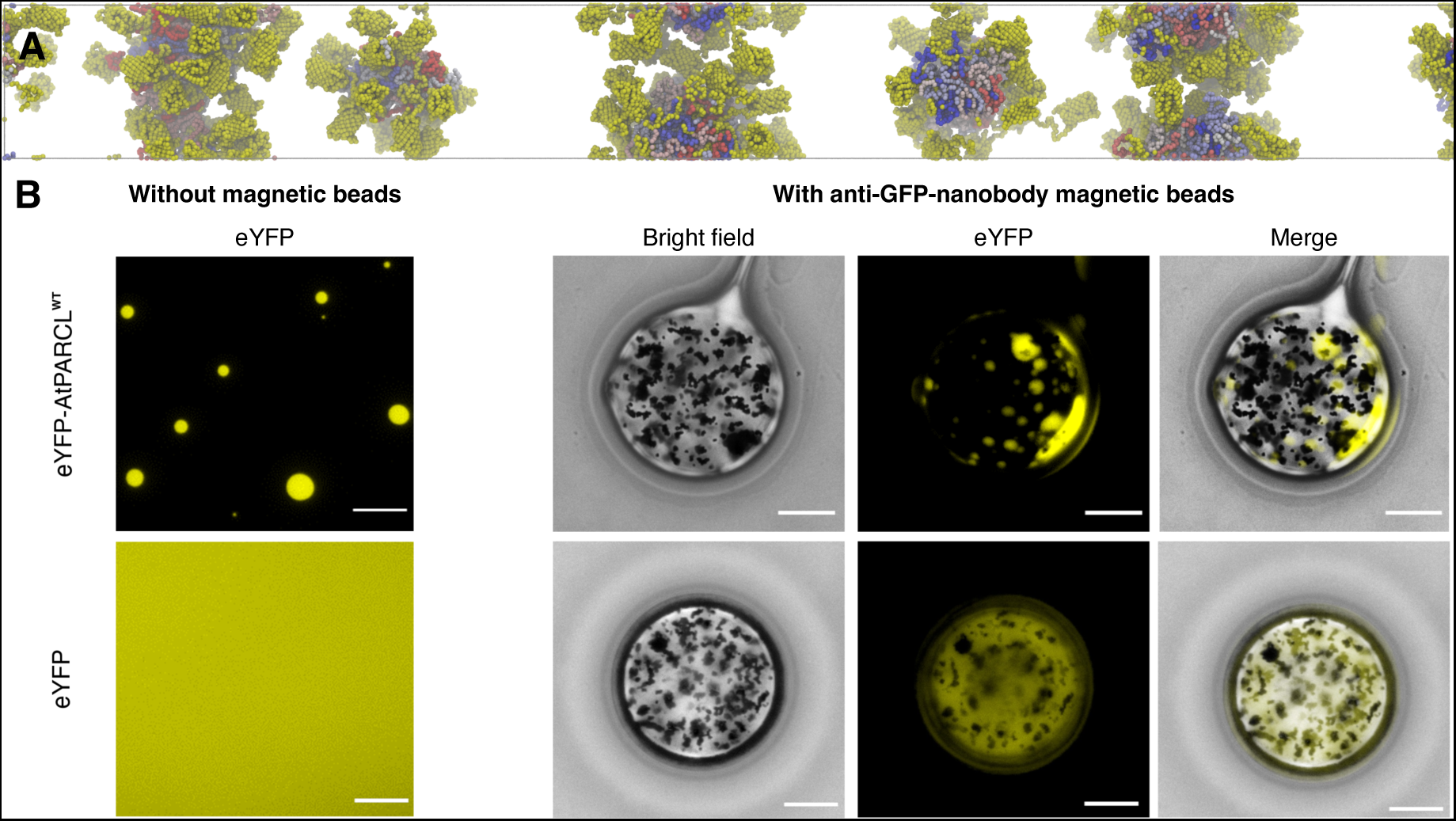
eYFP-PARCL phase separates with eYFP at the surface of the condensates. A Rendering of the coordinates of 120 eYFP-PARCL proteins after 100 nanoseconds of coarse-grained MD simulation. The eYFP domains are coloured yellow and treated as structured; each molecule’s PARCL domain is assigned a unique shade of the blue-white-red spectrum and treated as intrinsically disordered. Cluster formation is indicative of phase separation. The eYFP domains tend to reside at the cluster surfaces. Rendered in VMD using the Tachyon ray-tracer. B Testing eYFP accessibility in eYFP-PARCL condensates *in vitro* using magnetic beads coupled to anti-GFP-nanobodies. 20 µM protein was used and condensation was induced by adding 10 % PEG3350 in 1x condensation buffer. In contrast to free eYFP not undergoing LLPS, which shows uniform binding to the magnetic beads, eYFP-tagged PARCL condensates can be isolated on the beads, supporting the simulation results of solvent exposed eYFP in the eYFP-PARCL condensates. Scale bar: 10 µm.

### Phase separation of eYFP-PARCL can be attributed to the low-complexity domain of PARCL

Although eYFP alone was unable to phase separate under the conditions set (Fig 2B and Fig 3), the addition of a large tag to a given protein might inadvertently change the overall phase separation propensity of the fusion construct compared to the untagged protein. Despite using identical molar concentrations in all assays and simulations, a duplication of the molecular size will at least increase potential crowding effects as long as the overall reaction volume and conditions are kept constant, potentially influencing condensate size, number, or fluidity when compared to untagged proteins. To investigate the impact of eYFP on phase separation, we conducted further simulations using eYFP and PARCL alone. The FuzDrop (Hardenberg et al, 2020) LLPS predictor does not predict that eYFP undergoes phase separation; its score (pLLPS) of 0.4443 falls below the 0.5 threshold. In line with this, the MD simulations of eYFP show an approximately uniform distribution of molecules (Fig 3A). We next carried out MD simulations for PARCL (pLLPS=0.9969) without the eYFP tag. These simulations exhibit clear phase separation behaviour (Fig 3B) with PARCL forming three main droplets with some individual proteins remaining in the bulk solvent. Finally, we simulated the PARCL and eYFP proteins as separate molecules in the same system. Fig 3C shows the results using default parameters, and Fig 3D shows the same system after simulating with the altered parameters chosen for the eYFP-PARCL simulations. In both cases the PARCL proteins formed condensed droplets while the eYFP remained in the bulk solvent. *In vitro* experiments confirmed that eYFP alone does not form condensates, but that eYFP-PARCL does *in vitro* and *in vivo* (Fig 3E-G). Consistent with the simulations (Fig 3C-D), added free eYFP did not interfere with PARCL condensate formation and was excluded from condensates (Fig 3E).

**Figure 3.**
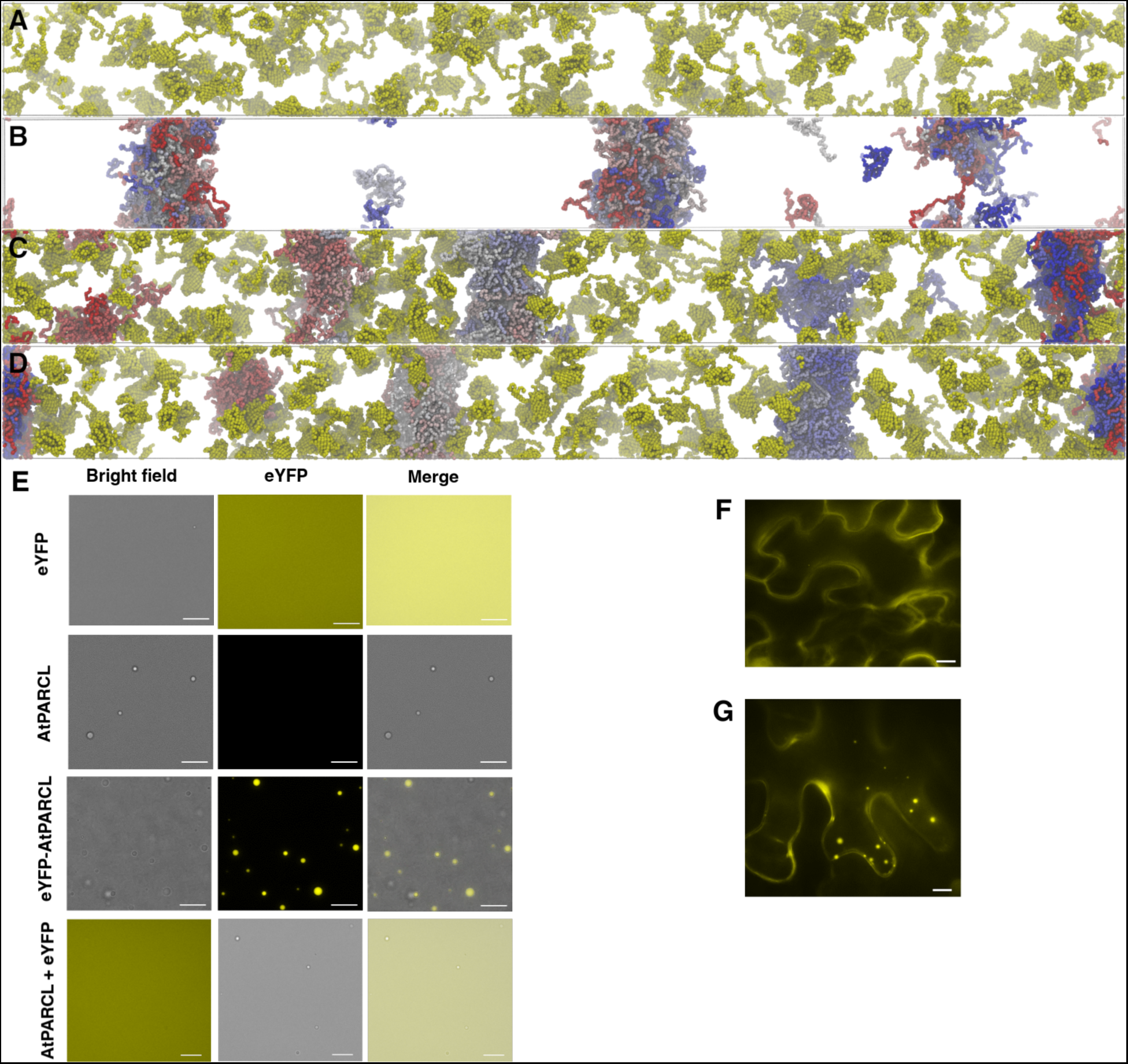
Condensation of eYFP-PARCL is driven by the PARCL domain rather than the eYFP domain. A Slab simulation of 120 structured eYFP proteins after 100 nanoseconds of coarse-grained MD simulation. No phase separation can be observed. Rendered in VMD using the Tachyon ray-tracer. B Slab simulation of 120 disordered PARCL proteins after 100 nanoseconds of coarse-grained MD simulation. The majority of the PARCL proteins are located in three dense “droplets”, with some isolated molecules and smaller clusters remaining in the dilute phase. Rendered in VMD using the Tachyon ray-tracer. C Slab simulation of 120 free eYFP proteins (yellow) and 120 wild-type PARCL proteins (colours along the blue-white-red spectrum), using default interaction and energy parameters. The PARCL proteins are clustered in 5 “droplets” which do not recruit the eYFP molecules; compared to the PARCL proteins alone in (B), there are more, smaller droplets shown in (C), and fewer isolated PARCL proteins. Rendered in VMD using the Tachyon ray-tracer. D 120 free eYFP proteins (yellow) and 120 wild-type PARCL proteins (colours along the blue-white-red spectrum), with energy and interaction parameters adjusted to accommodate the size of the eYFP domain. The PARCL proteins form 4 “droplets” with no isolates, while the eYFP remains in the bulk solvent. Rendered in VMD using the Tachyon ray-tracer. E Experimental validation of simulation result from A to D. Proteins were tested for their condensation behaviour after addition of 10 % PEG3350: free eYFP alone; free AtPARCL; eYFP fused to AtPARCL; an equimolar mixture of free eYFP and free AtPARCL.. Whereas free eYFP did not show any phase separation, free AtPARCL and a mixture of free eYFP and AtPARCL did and were in good agreement with the obtained simulation results. All proteins were measured at 10 µM concentration in 1x condensation buffer and 10 % PEG3350 with the same exposure time (17 ms). Pictures were taken using bright field and YFP fluorescence. Scale bar: 10 µm. F *In vivo* observation of free eYFP in agroinfiltrated tobacco leaves. The transiently expressed free eYFP did not show any condensates. Scale bar: 10 µm. G *In vivo* observation of PARCL condensates in agroinfiltrated tobacco leaves. Wild-type PARCL N-terminally fused to eYFP was transiently expressed in tobacco leaves. EYFP-PARCL showed several cytosolic condensates throughout the cells, whereas free eYFP (F) did not show any condensates. Scale bar: 10 µm.

These analyses suggest that whilst eYFP-PARCL forms condensates, this behaviour is due to PARCL’s disordered region and that without eYFP the tendency to phase separate would be even greater. This can be visualised in network graphs based on the intermolecular interactions. In this network, each protein is represented by a node; two nodes are connected if any residues from each molecule are within contact difference in the final frame. Communities of connected nodes indicate clusters or droplets, while unconnected nodes show isolated proteins. Fig 4 shows the differences in networks for PARCL with and without eYFP. Without the eYFP tag (Fig 4A), the proteins formed two large droplets and one smaller droplet during the simulation, and eight proteins were not inside any of the droplets in the final frame. With eYFP tagging (Fig 4B) the final frame of the trajectory showed 6 smaller droplets with no isolated proteins. Fig 4C shows the result with equal amounts of free eYFP, which showed the same increased separation as Fig 4B.

**Figure 4.**
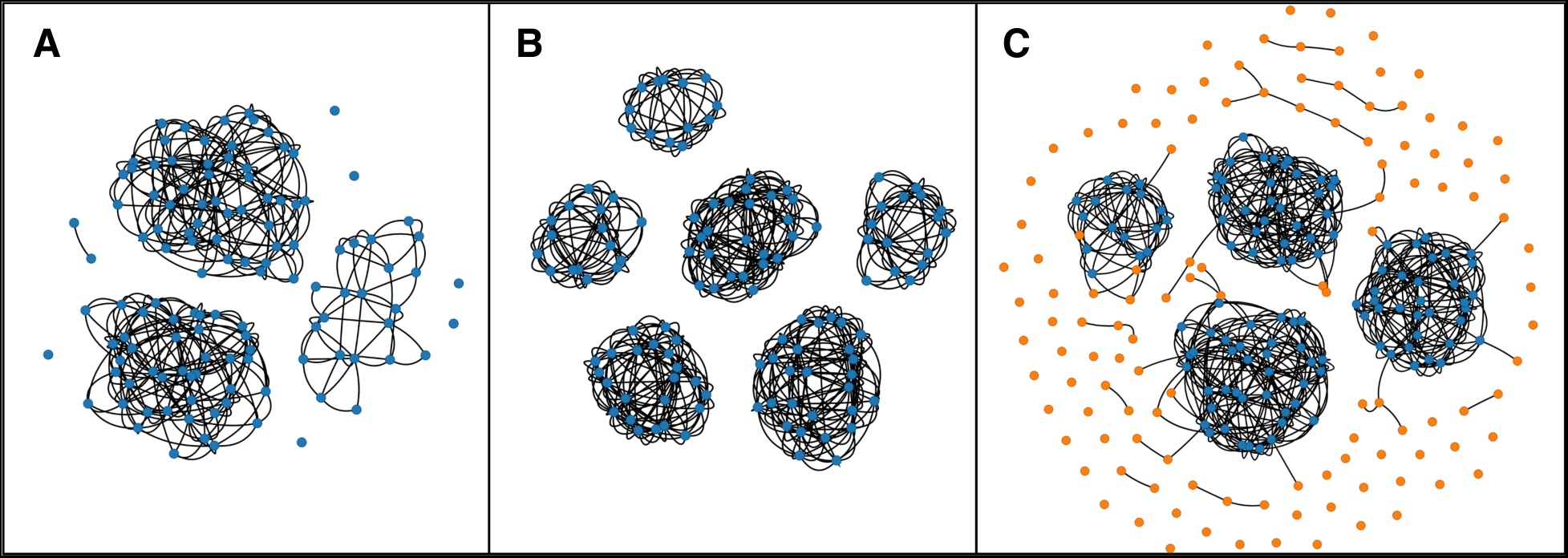
The inclusion of eYFP results in more droplets which are smaller and more tightly clustered. Network graph representation of the final frames of coarse-grained MD simulation of eYFP-PARCL. Each node represents a protein, and edges represent that at least one pair of residues from the node proteins are within contact distance. A Simulation of PARCL^WT^. The 120 nodes are 120 wild-type PARCL proteins. The nodes form three well-connected communities which appear as droplets in the rendered images, with some isolated nodes that represent PARCL proteins remaining in the dilute phase. B Simulation of eYFP-PARCL^WT^. The 120 nodes are 120 eYFP-PARCL proteins. The nodes form 6 tightly-connected communities, which are represented as clusters or small droplets in the rendered images. All nodes are included in one of these communities; there are no isolated proteins in the dilute phase. C Simulation of PARCL^WT^ with added free eYFP. The 120 blue nodes are 120 wild-type PARCL proteins and the 120 orange nodes are 120 free eYFP proteins. The PARCL nodes form 4 communities, which are represented as droplets in the rendered images. The eYFP nodes are mostly isolated, although some have a small number of neighbours. This is likely to be caused by the very high concentration of molecules within the system.

Despite the overall behaviour of PARCL and eYFP being consistent in terms of condensate formation, some differences could be identified. Visible observation of the simulations showed that the droplets of eYFP-PARCL grew slower than those of PARCL alone. Small clusters that meet merged less readily when the eYFP domain is present than without. This may be due to the way the PLD, with its interacting aromatic residues, tends to be buried in the centre of the clusters, leaving the eYFP domain at the interface. This results in PLDs that are less accessible to molecules outside of the droplet. Additionally, the eYFP domains may pose an obstacle to proteins that would otherwise diffuse in and out of the droplet, resulting in clusters with more consistent membership.

From these results, we conclude that coarse-grained MD simulations can successfully reproduce condensate formation of the PARCL protein (with and without eYFP) and that PARCL alone is sufficient to induce phase separation, in line with *in vitro* studies (Fig 3E). We find that whilst the inclusion of eYFP makes the simulations more challenging (and significantly more time consuming), we can reproduce the experimental results and simulate condensate formation of eYFP-PARCL and the deviation from using untagged PARCL is minimal. In the interest of compute time, for most of the simulations we can therefore use only PARCL and then validate against experimental data and simulations using eYFP-PARCL can be minimised to selected cases.

Having gained confidence in the ability to reproduce experimental observations of condensate formation *in silico*, we wanted to ascertain the driving factors for this phenomenon. Following previous suggestions that macromolecule:macromolecule interactions and multi-valency are important for phase separation, we reasoned that those residues that come into contact with other molecules might play a role in PARCL condensate formation. To estimate contact frequencies, we used the molecular dynamics trajectories to identify pairs of residues that pass within contact distance, as calculated from the radii of each amino acid. Analysing these potential interactions between residues of different PARCL molecules allowed us to determine which parts of the protein are contributing to the phase separation behaviour observed in the simulation. Figure 5 shows how many intermolecular contacts each of the PARCL residues make, averaged across the simulation. Aromatic residues, particularly tyrosine, have been shown to participate in interactions that promote LLPS (Das et al, 2020; Martin et al, 2020). In agreement with this, the simulations showed that, apart from the leading methionine, PARCL’s 10 tyrosines make the most intermolecular contacts. The first four tyrosines are located in pairs at the start of the sequence, and the remaining six appear throughout the protein’s PLD. The four paired residues were the least-interacting tyrosines. To confirm that the condensed phase was liquid-like we compared the contacts between frames, looking at how often contacts are made and broken. In line with the nature of intermolecular interactions expected of liquids, we found that the contacts were transient: In the untagged PARCL simulation, 99.7% of the residue pairs that came within contact distance in a frame of a trajectory had separated by the next frame, 100 picoseconds later. With the eYFP tag present this was reduced to 99.6% of contacts. This supports our visual assessment that the droplets were highly dynamic and fluid-like.

**Figure 5.**
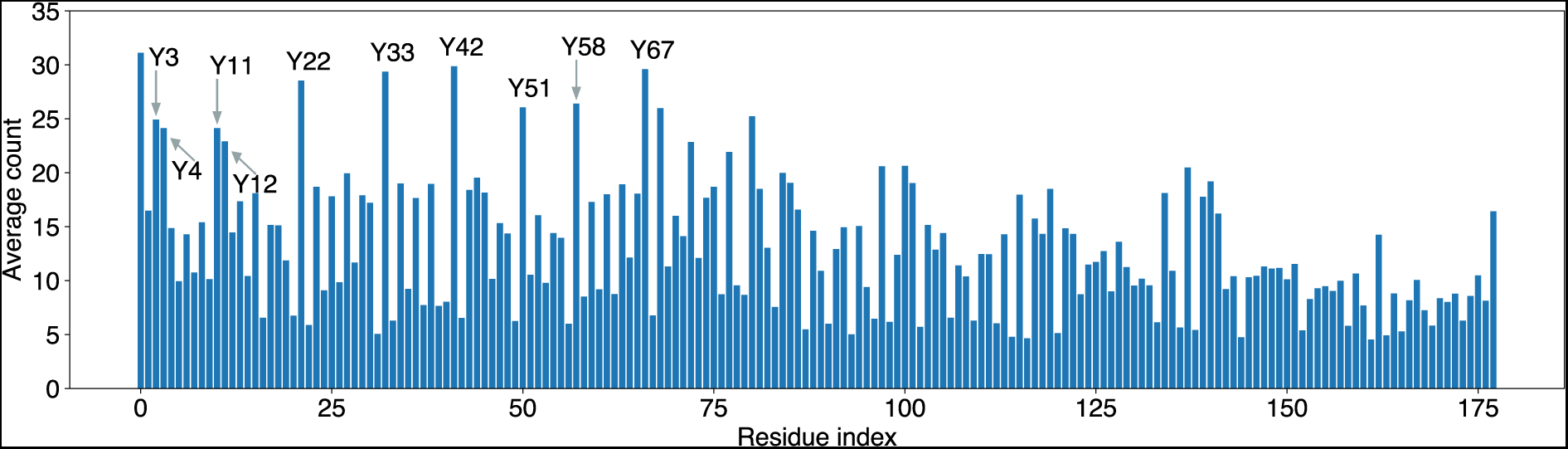
Potential contact frequency between residues determines key drivers for phase separation. Average number of potential contacts per frame for the 120 copies of each residue of the wild-type PARCL protein during 1000 frames of coarse-grained MD simulation, lasting 100 nanoseconds. Two residues from different PARCL molecules are considered in potential contact if the distance between the coordinates of their representative beads is less than the sum of their radii. While potential contacts were spread across the whole PARCL sequence, there were more noticeable peaks in its N-terminal half. Many of the peaks most often falling within contact distance are tyrosine residues, which are labelled by residue index.

The key interactions in the protein can be visualised in a network, Supplementary Figure 2, where nodes represent residues and edges indicate contacts between the residues. Many of the residues encountered one another across the entire trajectory (for clarity Supplementary Figure 2 shows only interactions that occurred in at least 300 frames). The aromatic residues of the PLD formed a large, well-connected cluster in the network through frequent interactions with one another as well as contacts with other residues, particularly leucine residues adjacent to the PLD. The residues at each end of the sequence were in the subset of high-frequency contact residues, suggesting a potential role in phase separation. The initial methionine residue was included in the PLD cluster, and the final aspartic acid made frequent contact with several residues in a short region towards the C terminal that were not highly contacted individually.

To evaluate the impact of eYFP on the interacting residues, we conducted the same analysis using eYFP-PARCL. We counted contacts from the trajectories to determine whether eYFP introduced changes in the intermolecular interactions of PARCL. We found that despite the inclusion of the eYFP domain, the general ranking of the PARCL residues by contact frequency remained the same, apart from a reduced role for the initial methionine which was hindered by its adjacency to the eYFP structure. The residues of eYFP themselves made only minimal contact to other molecules. The PLD tyrosine residues that were identified as important in Figure 5 were again the most contacted residues in the eYFP-PARCL simulations (Supplementary Figure 1).

To summarise, the analysis of intermolecular residue contact frequencies can be used to pinpoint potential key residues in the protein. Furthermore, the impact of eYFP on the predicted interactions was minimal, affecting only residues adjacent to the eYFP domain.

### Targeted engineering of PARCL validates the predicted phase separation behaviour

Having shown that the simulations using eYFP-PARCL can capture the experimental system, we proceeded to evaluate whether the simulations could be used for rational, targeted perturbations to condensate formation. Using the simulations’ contact frequencies as a guide, we set up simulations with an *in silico* PARCL mutant that lacked the residues that appeared to be driving the phase separation behaviour. As the most frequently contacted positions, the 6 tyrosine residues of the PLD were replaced with glutamic acid (PARCL^PLD^ ^Y-E^), since phosphorylation studies on wild-type PARCL highlighted that tyrosine phosphorylation at these sites can prevent condensation (Ostendorp 2022). Fig 6A and Fig 6B show renders of the final frames of the MD simulations, without fluorescent tags and with eYFP, respectively. In the eYFP-PARCL simulation (Fig 3B,), there were some small clusters made up of the PARCL regions, but Fig 3A showed little evidence of LLPS. We have plotted the contact networks in Fig 6C and Fig 6D, which shows the connections between the clusters. To test these predictions and experimentally validate phase separation behaviour of mutants of the residues that the MD simulations had highlighted as being potentially important interactions, we tested experimentally if PARCL^PLD^ ^Y-E^ proteins can form condensates *in vitro* and *in vivo* in transiently overexpressing tobacco leaves. In line with simulations, PARCL^PLD^ ^Y-E^ was unable to form any condensates (Fig 6E and F).

**Figure 6.**
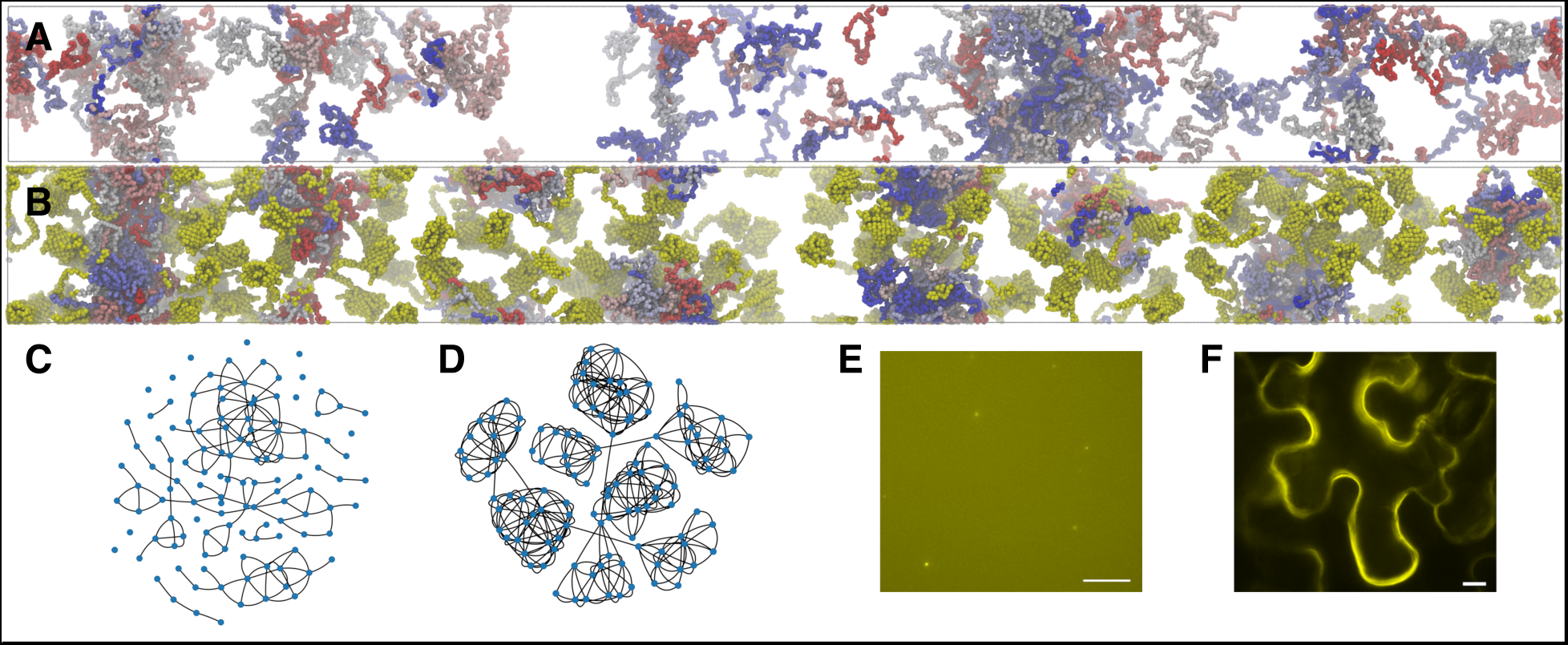
Mutating residues with high estimated contact frequency results in diminished phase separation. The 6 tyrosine residues that were most often potentially contacted in the simulation of the wild-type (at indices 22, 33, 42, 51, 59, and 67) have been replaced with glutamic acid. A Rendering of the final frame of the 100 nanosecond coarse-grained slab simulations performed on the mutated PARCL^PLD^ ^Y-E^ protein. While there are some clusters, they appear looser and less well-defined than the dense droplets observed in the wild-type simulations. B Rendering of the final frame of the eYFP-PARCL^PLD^ ^Y-E^ mutant after 100 nanoseconds of coarse-grained MD simulation. There are some small clusters, where the PARCL domains are making apparent contact with one another and the eYFP domains remain at the interface. There are molecules distributed across the slab’s long *z*-axis, without clear regions of solely bulk solvent as seen in the wild-type simulations. The clusters here are more indicative of phase separation than in the untagged PARCL simulation (B), which we attribute to the crowding effect of the eYFP domain on the already highly concentration of the system. C Network graph representation of the final frame of the PARCL^PLD^ ^Y-E^ MD simulation that is rendered in (A). Each node represents a protein, and edges represent that at least one pair of residues from the node proteins are within contact distance. The number of isolates and very small communities has increased from the wild-type simulation, and the clusters are now more loosely connected. D Network graphs showing the final frame of the eYFP-PARCL^PLD^ ^Y-E^ MD simulations that is rendered in (B). Each node represents a protein, and edges represent that at least one pair of residues from the node proteins are within contact distance. The presence of the eYFP domain has resulted in many small clusters which occasionally come into contact distance with each other, shown by the communities in the graph. E *In vitro* condensation of PARCL^PLD^ ^Y-E^ mutants. 10 µM protein was used and condensation was induced by adding 10 % PEG3350. Scale bar: 10 µm. F EYFP-PARCL^PLD^ ^Y-E^ *in vivo* condensation in agroinfiltrated tobacco leaves. In line with simulations and *in vitro* assays, PARCL^PLD^ ^Y-E^ was unable to form condensates in cells. Scale bar: 10 µm.

This allowed us to compare the changes in phase separation *in vitro* and *in silico*. Fig 6E shows the *in vitro* results of using a PARCL mutant with 6 tyrosine residues in the PLD mutated to glutamic acid. In the experiments we included crowding agents to achieve the required concentration for *in vitro* phase separation. In the MD simulations, we do not mimic the exact concentration of PARCL and PEG, instead we use a high concentration of PARCL molecules to account for the crowding effect of PEG. By adding the eYFP’s extra length to the proteins we have increased the eYFP-PARCL concentration to an even higher level than that of the experiment, causing a difference in phase separation behaviour. The clusters could also reflect what is happening in the *in vitro* experiments as while the PARCL sections of the molecules are interacting in small clusters, the eYFP domains are broadly evenly concentrated throughout the slab.

To summarise, we have shown the utility of intermolecular contact maps by making predictions for phase separation behaviour that we could reproduce both *in silico*, in *vitro and in vivo*. This methodology may be of general applicability for predicting target residues for experimental perturbation.

### Alternate tyrosine mutations show importance of residue selection

The PARCL^PLD^ ^Y-E^ mutant was created by mutating the residues (tyrosines) that made the most frequent contacts in simulations of PARCL^WT^. We varied the tyrosines chosen for mutation to compare the PARCL^PLD^ ^Y-E^ mutant’s LLPS disruption with other possible choices of tyrosine residues to mutate. From an initial set of wild-type PARCL coordinate and input files, we programmatically replaced alternative combinations of tyrosines with glutamic acid for MD simulation. Based on our initial work each simulation was run for 100 nanoseconds.

We began with one simulation in which all tyrosines were replaced, ten simulations where each of the ten tyrosines was mutated alone, and 20 simulations where random combinations were chosen. For the combinations, four sets of two indices were chosen at random, followed by four sets of three indices, and continuing up to four sets of six indices. Not all possible combinations of residues were chosen for each set size due to the computational expense of evaluating all of the hundreds of possible sets. We also performed the simulations with a smaller subset of the PLD tyrosines, leaving out the tyrosines at indices 51 and 58, which appeared to be involved in fewer contacts as shown in Figure 5, and another simulation using only the 5 tyrosine residues in the PLD. For one simulation, we mutated the four tyrosines outside of the PLD at indices 3, 4, 11, and 12, which were not changed as part of the original mutation experiment.

We summarised the LLPS behaviour for each mutant as a single metric for ease of comparison between simulations by using the IDR HPS energy reported by GENESIS for each frame of the simulation. Fluctuation around a negative number indicates the formation of a condensate, whereas numbers closer to zero indicate a lack of LLPS. We average across the final 50 frames to give a smoother representation of the converging point of each simulation. Fig 7 shows the mean IDR HPS energy in this period for each of the MD simulations. The number of mutated residues appeared to be roughly negatively correlated with the HPS energy, but the choice of which residues to mutate has a large effect. For small numbers of mutations, choosing tyrosines that were most frequently interacting in the wild-type simulations does not have the most destructive effect on phase separation according to the HPS energy. In the case of the residue at index 22, mutating only a very frequently interacting residue has resulted in simulations that indicate a marginally stronger phase separation. For larger groups of mutated residues, phase separation was more diminished when the tyrosines that were replaced frequently came into contact in the original simulations.

**Figure 7:**
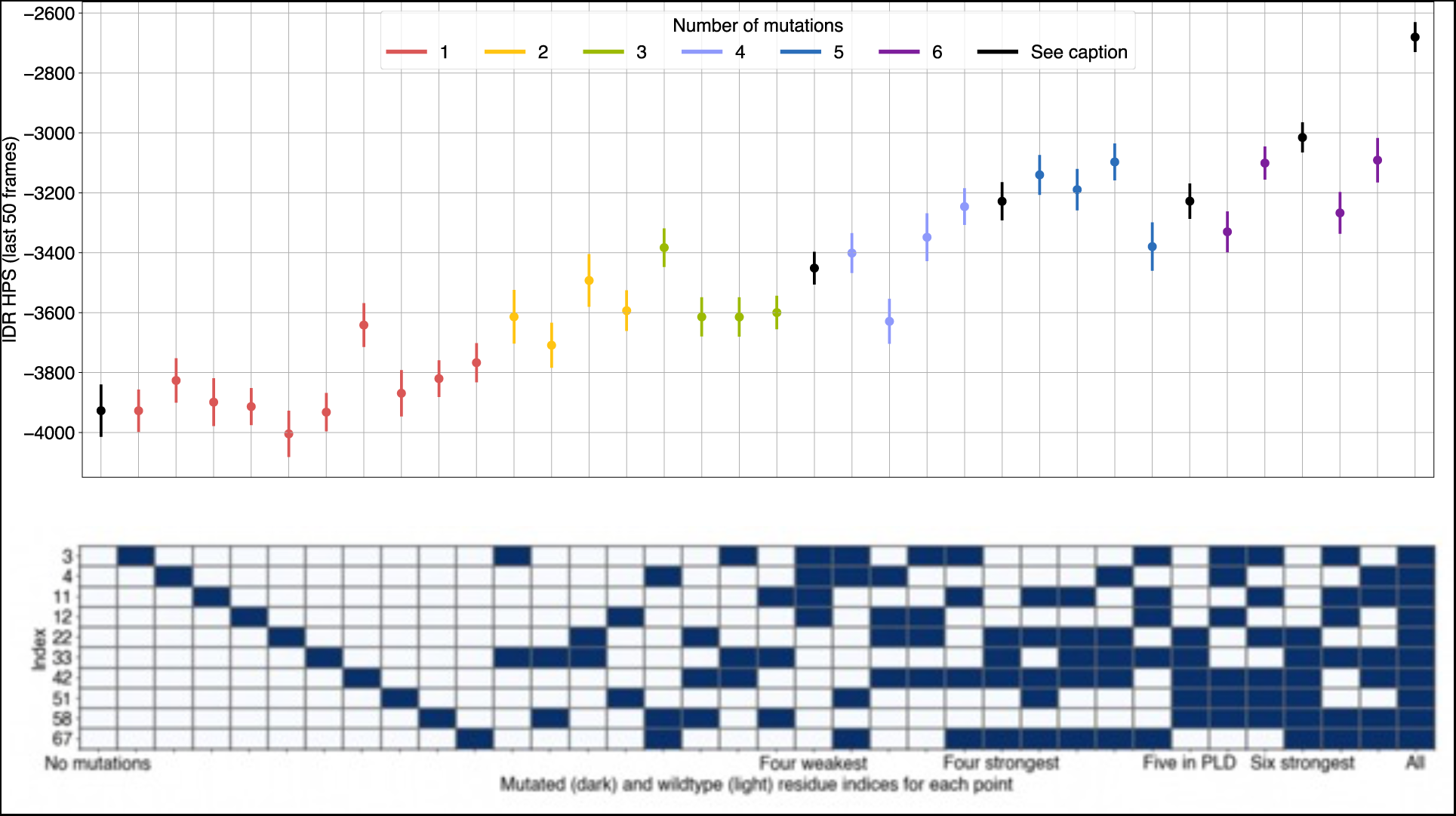
Broadly, mutating more tyrosines results in less pronounced LLPS. However, choice of specific residues can be highly influential, which the model is able to reflect in these simulations. The PARCL^PLD^ ^Y-E^ mutant (designed from the PARCL^WT^ simulation results) yielded the second-highest disruption, second only to the mutant in which all were replaced. In some cases the result is predictable from contact distance analysis of the wild-type simulation, but in others the apparent influence of residues is unexpected, indicating that the phase separation propensity of a sequence is not simply a result of a linear combination of individual residues’ contribution. Points show mean IDR HPS energy contribution reported by the GENESIS simulation log files between frames 1,950 and 2,000 of the slab simulation trajectories with various mutations; lines show standard deviation.

PARCL’s well-spaced tyrosine residues offer a long stretch of sequence that appears to be favourable for the multivalent interactions characteristic to phase separation. Our simulations show that these residues are promiscuous in regard to partner residues, interacting with one another and with other residues, so individual mutations of the tyrosine residues are not enough to prevent phase separation. MD simulations like these can capture the effects of fine-grained mutation combinatorics.

### Simulations of miRNA-PARCL interactions align with experiments in both the RNA interaction site and phase separation behaviour

PARCL is known to bind miRNAs (Ostendorp, 2022). To evaluate whether this has an impact on condensate formation, we added miR399 to the simulations to determine how phase separation might occur in this two-component system.

### Simulations indicate a possible binding site containing most potential contacts between PARCL and miR399

When miR399 was included in the simulations, PARCL condensate formation occurred much as before, where the tyrosines in the PLD were residues that most frequently took part in PARCL-PARCL interactions and the PLD tended to form the centre of the resulting droplets. PARCL-miR399 interactions, as shown in Fig 8, rarely occurred in PARCL’s PLD, instead being heavily focused on a short region towards the C-terminus. The high-contact region in Fig 10 covers the arginine, histidine and lysines around residue index 150. These residues are not themselves interaction hubs in the PARCL-only simulations but, as shown in Fig 5, they are each frequently contacted by the final aspartic acid. This K-rich region was indicated by Ostendorp et al (2022) as a site for the binding of nucleic acids.

**Figure 8.**
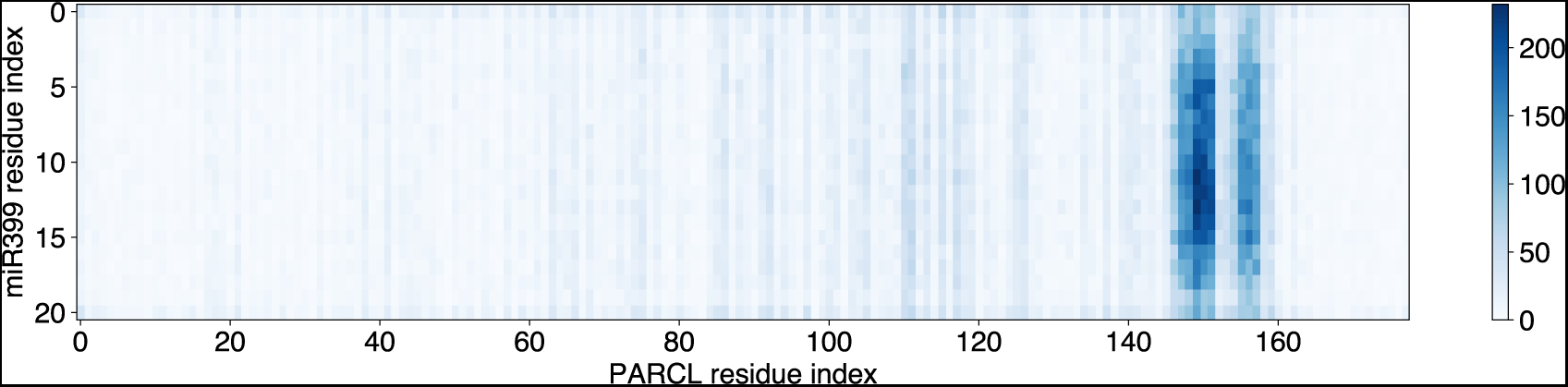
Potential contacts between PARCL and miRNA are mostly limited to a short, K-rich region towards the C-terminus. The heat-map shows the number and location of potential contacts between residues in PARCL and in miR399 in a coarse-grained MD simulation.

Potential contacts are counted when a residue from a PARCL protein and a residue from miR399 are closer together than the sum of their standard radii. The count is summed from 1000 frames (100 nanoseconds) of simulation. Potential contacts occur across the miR399 sequence, with a small preference for the central nucleotides. The PARCL proteins’ potential points of contact with the miRNA are largely limited to a short region towards the C-terminus rich in arginine, histidine, and lysine residues.

### Serine mutation in C-terminal domain has little effect on condensation

Ostendorp et al (2022) showed that phase separation of the PLD and RNA binding of the C-terminal region function independently and can be manipulated individually. Our simulations also pinpoint different residues driving condensate formation and nucleic acid binding.

Ostendorp et al (2022) mutated five serines in the C-terminal region to alanine residues and reported that phase separation still occurred, but RNA binding was reduced. We therefore repeated our PARCL simulations with these five serine residues mutated into glutamic acid. Fig 9A and 9B shows the results of this simulation, which were similar to the results of the wild-type simulation, with and without the eYFP domain, respectively. Fig 9D and 9E show the same frames represented as network graphs.

**Figure 9.**
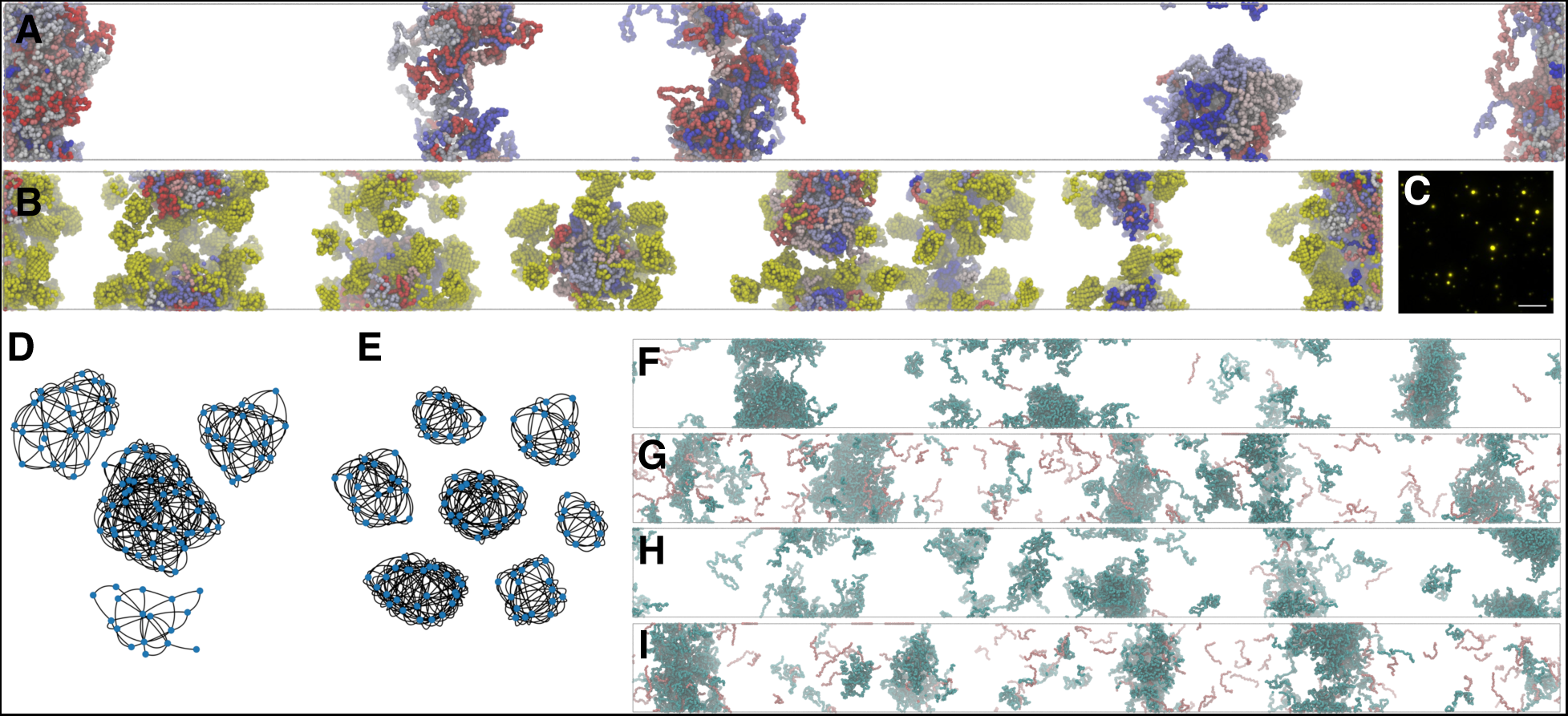
– Mutations that reduce RNA binding do not substantially affect LLPS. Additionally, phase separation of the wild-type and PARCL^C-term^ ^S-E^ mutant still occurs in the presence of miRNA, but simulations suggest smaller droplets. Serine residues near the C-terminus (at indices 169, 171, 172, 173, 175, and 177) have been replaced in the mutant. A Rendering of the final frame of the 100 nanosecond coarse-grained slab simulations performed on the mutated PARCL protein. All protein molecules are contained within four dense droplets, indicating phase separation as in the simulations of wild-type PARCL. B Rendering of the final frame of the 100 nanosecond coarse-grained slab simulations performed on the eYFP-PARCL^C-term^ ^S-E^ mutant. Cluster formation is indicative of phase separation. The eYFP domains remain at the cluster interfaces. C *In vitro* condensation of PARCL ^C-term^ ^S-E^ mutant. 10 µM protein was used and condensation was induced by adding 10 % PEG3350. Scale bar: 10 µm. Small droplets are clearly visible. D Network graph representation of the final frame of the mutated PARCL MD simulation that is rendered in (A). Each node represents a protein, and edges represent that at least one pair of residues from the node proteins are within contact distance. The four well-connected communities are similar to the three communities observed in the wild-type simulation, but there are no isolated PARCL molecules. E A network graph showing the final frame of the MD simulation of the eYFP-PARCL^C-term S-E^ mutant. Each node represents a protein, and edges represent that at least one pair of residues from the node proteins are within contact distance. As in the wild-type simulations, the addition of the eYFP domain has resulted in more, smaller communities. F Rendering of the final frame of the 100 nanosecond coarse-grained slab simulations performed on 120 mutated PARCL proteins (cyan) and 12 miR399 (red) present. The miR399 molecules were primarily in and around the PARCL droplets, and the droplets appeared smaller, with more PARCL proteins that are isolated or in small clusters with miRNA. G Rendering of the final frame of the 100 nanosecond coarse-grained slab simulations performed on 120 mutated PARCL^C-term^ ^S-E^ proteins (cyan) and 120 miR399 (red) present. There are miRNAs within and at the interface of the condensates, but also isolated throughout the bulk solvent, which could suggest a limit to the amount of miR399 that can be recruited by droplets of these sizes. H Rendering of the final frame of the 100 nanosecond coarse-grained slab simulations performed on 120 PARCL^WT^ proteins (cyan) and 12 miR399 (red) present. As in (F), the miR399 molecules are primarily in and around the PARCL droplets. The droplets are smaller, more abundant, and less well-defined than simulations without miRNA. I Rendering of the final frame of the 100 nanosecond coarse-grained slab simulations performed on 120 PARCL^WT^ proteins (cyan) and 120 miR399 (red) present. There are miRNA within and at the interface of the condensates, but also isolated throughout the bulk solvent, which could suggest a limit to the amount of miR399 that can be recruited by droplets of these sizes.

We characterised each system’s interactions by calculating how often the different types of molecules came within contact distance (Table 1). Both the PARCL^WT^ and PARCL^C-term S-E^ proteins interacted with the miRNA, but the mutant tended to interact more with other PARCL proteins and less with the miR399 molecules than the PARCL^WT^. Fig 9 F-I show the disruptive effects the presence of RNA has on the phase separation at these concentrations, which matches the smaller droplets seen in experiments (Fig 9C).

**Table 1:** The C-terminal mutant Y-E increases PARCL-PARCL interactions but reduces interactions between PARCL and miR399.

| System composition | Average interactions per frame of trajectory |  |  |
| --- | --- | --- | --- |
|  | PARCL-PARCL | PARCL-miR399 | miR399-miR399 |
| 120 wild-type PARCL<sup>WT</sup>, 12 miR399 | 776.23 | 10.35 | 0.00 |
| 120 wild-type PARCL<sup>WT</sup>, 120 miR399 | 559.36 | 65.57 | 0.02 |
| 120 PARCL<sup>C-term S-E</sup> mutant,<br>12 miR399 | 935.97 | 3.84 | 0.00 |
| 120 PARCL<sup>C-term S-E</sup> mutant,<br>120 miR399 | 883.86 | 33.61 | 0.01 |

The results of this section demonstrate that miRNA binding is likely independent from the regions of PARCL that drive phase separation. Again, the simulations are consistent with the experimental data.

In summary, our analysis has shown that coarse-grained MD simulations coupled with an analysis of contact dynamics can be used as an effective tool to make predictions about phase separation behaviour and to guide targeted and rational engineering of proteins to alter condensate formation.

## Discussion

Liquid-liquid phase separation has recently been identified as an important mechanism in biology for compartmentalising cells. This process leads to the formation of biomolecular condensates that have been implicated in stress granules, transcription machinery and protein degradation. Our understanding of the nature of condensate formation has advanced significantly through experimental and computational approaches, including the development of powerful molecular dynamics frameworks to study phase separation *in silico.* Here, we applied a coarse-grained MD model to guide predictive protein engineering approaches. We studied the plant protein PARCL that has been shown to exhibit condensate formation *in vivo* and *in vitro*. We use this protein as a case study for evaluating the agreement between our framework of computational predictions and experiments (Fig 1), and to explore the predictive power for the rational design of targeted mutations.

We performed coarse-grained MD simulations of the phloem protein PARCL using the hydrophobicity scale model proposed in Regy et al (2021), allowing us to simulate, visualise and quantify the protein’s phase separation behaviour (Fig 2A). Phase separation of PARCL can be observed experimentally *in vitro* and *in vivo* using fluorescent tagging (Ostendorp 2022). These tags have previously been elided from MD simulations, but have the potential to affect molecules’ behaviour where, as in the case of PARCL, they are a larger folded domain appended to an otherwise intrinsically disordered protein. We therefore carried out MD simulations with eYFP-PARCL proteins for a more accurate reflection of the experiments. We were able to reproduce phase separation with the eYFP-tag, using changes to the model’s parameters that did not affect the results of our control simulations (Fig 3C-D). The resulting condensate droplets were each structured with the PARCL domain within and the eYFP domain at the phase boundary. While we can’t exclude this being an artefact of the forcefield and model, we provide experimental data supporting this structural arrangement, as the eYFP domain was accessible to anti-GFP-nanobodies coupled to magnetic beads (Fig 2B). eYFP-PARCL forms more droplets than PARCL, and these droplets are smaller and more tightly-connected (Fig 4A-B). Similar droplet networks occur if free eYFP is added to PARCL simulations (Fig FC). We confirmed that the eYFP domain is not driving the LLPS as it does not form condensates alone *in silico* (Fig 3A), *in vitro* (Fig 3E), or *in vivo* (Fig 3F). eYFP-PARCL does form condensates *in silico* (Fig 2A), *in vitro* (Fig 3E), and *in vivo* (Fig 3G). Amino acid contact frequency is similar in the tagged and untagged PARCL molecules (Fig EV1, Fig 5).

From a simple, radius-based analysis of which amino acids came within contact distance during the simulations, we identified candidate residues that may drive phase separation: a web of interactions centred around tyrosines (Fig 5, EV Fig 2) in the prion-like domain. While all the tyrosines within the PARCL sequence were involved in this network, this analysis highlighted which tyrosine residues had higher or lower involvement. We designed a mutant, PARCL^PLD^ ^Y-E^, in which the 6 tyrosines with the highest contact frequency were changed to glutamic acid. We confirmed that this mutant is less prone to phase separation *in silico*, *in vitro* and *in vivo* (Fig 6). In further simulations, we varied which of the tyrosines were mutated to glutamic acid, providing a wider scale of mutation testing *in silico* than would be practical in experiments. This showed how mutating different amounts of the tyrosines affects the phase separation behaviour (Fig 7). Comparing the strength of the effect showed that while reducing the number of tyrosines generally reduced phase separation, the choice of which residues to mutate is more important. For each quantity of mutated residues, there was a significant range of HPS energy scores. In some cases, this followed the expectations established by the PARCL^WT^ simulations, such as the simulation removing the four most potentially contacted tyrosine residues which showed the most effective disruption of LLPS of any combination of four. In other cases, this differs from expectations, such as the four least contacted tyrosine residues; a random selection of four tyrosines had less effect on LLPS as indicated by the HPS energy. Our simulations showed that the PARCL^PLD^ ^Y-E^ mutant was more disruptive to LLPS than any other tested mutant with 6 or fewer mutated residues. The simulations also demonstrated that phase separation behaviour cannot be fully predicted from a linear combination of the contribution of individual amino acids, but that the HPS-Urry model can simulate the effects of mutations. Developing a more sophisticated method of learning the effects of potential mutations or proposing mutations for a desired effect without performing exhaustive combinations of simulations would be an interesting strand to follow and is something we are exploring.

We included both miRNA and PARCL in a simulation and found that, even though the simulation did not attempt to accurately model the miRNA’s structure, it was capable of precisely identifying the protein’s nucleic acid binding region (Fig 8). We simulated a mutant, PARCL^C-term^ ^S-E^, in which 6 serine residues near the C-terminus were mutated to glutamic acid. These residues showed low contact propensity in wild-type simulations, and the mutants were able to phase separate with and without the presence of miRNA (Fig 9). The simulated mutants made less contact with miRNA than the wildtype, recapitulating experimental findings that phase separation and RNA binding can be independently targeted in PARCL (Ostendorp et al, 2022).

This study has demonstrated the effectiveness of coarse-grained MD simulations and contact analysis to guide mutational studies aimed at perturbing condensate formation and protein-RNA interaction. Moreover, all the *in silico* predictions could be experimentally validated *in vitro* and even *in vivo*. The introduction of well folded tagged proteins into simulations will help to increase their predictive power and can be used to examine the effects of fluorescent tagging on phase separating proteins. Simulations typically neglect folded domains or additional protein tags that are necessary for *in vivo* studies. This discrepancy might lead to contradictory results from simulations and *in vivo* observations. Our approach may help overcome this issue. We believe the workflow presented here may be of general applicability for targeted engineering of condensate formation and protein-RNA interactions.

## Acknowledgements

This article is part of a project that has received funding from the European Research Council (ERC) under the European Union’s Horizon 2020 research and innovation program (Grant agreement No. 810131), and supported by the Deutsche Forschungsgemeinschaft (KE 856/8-1 to J.K.).

## Conflict of Interest

The authors declare that they have no conflict of interest.

## Expanded View Figures

**Expanded View Figure 1:**
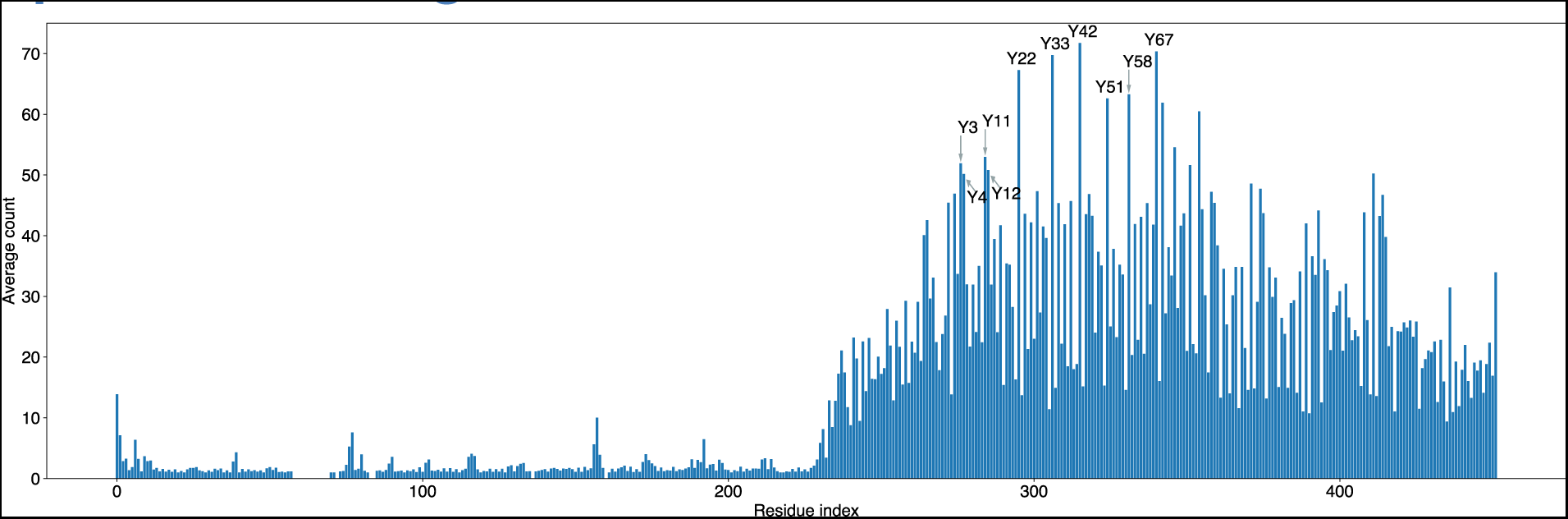
eYFP-PARCL residue average contact counts. The first 274 residues belong to the eYFP domain. Tyrosine residues have been labelled according to their position in the PARCL sequence.

**Expanded View Figure 2:**
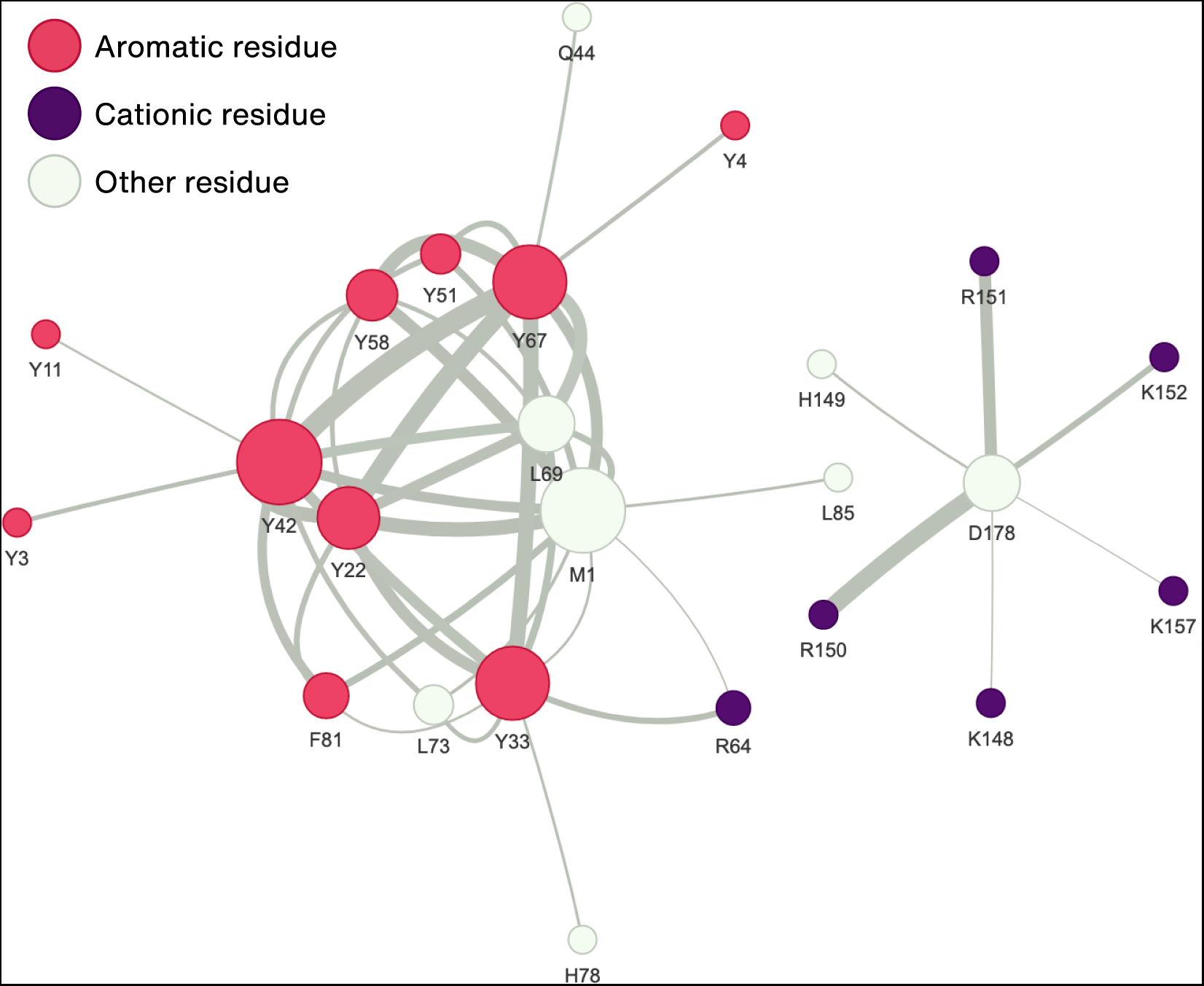
Residue pairs that most frequently interact in the PARCL wild-type simulation. Node size represents number of contacts between the residue and any other residue. Edge width represents number of contacts between the two connected nodes.

## Notes

### Competing Interest Statement

The authors have declared no competing interest.

